# Context-aware genomic surveillance reveals hidden transmission of a carbapenemase-producing *Klebsiella pneumoniae*

**DOI:** 10.1101/2021.06.07.447408

**Authors:** Adrian Viehweger, Christian Blumenscheit, Norman Lippmann, Kelly L. Wyres, Christian Brandt, Jörg B. Hans, Martin Hölzer, Luiz Irber, Sören Gatermann, Christoph Lübbert, Mathias Pletz, Kathryn E. Holt, Brigitte König

## Abstract

Genomic surveillance can inform effective public health responses to pathogen outbreaks. However, integration of non-local data is rarely done. We investigate two large hospital outbreaks of a carbapenemase-carrying *Klebsiella pneumoniae* strain in Germany and show the value of contextual data. By screening more than ten thousand genomes, 500 thousand metagenomes, and two culture collections using *in silico* and *in vitro* methods, we identify a total of 415 closely related genomes reported in 28 studies. We identify the relationship between the two outbreaks through time-dated phylogeny, including their respective origin. One of the outbreaks presents extensive hidden transmission, with descendant isolates only identified in other studies. We then leverage the genome collection from this meta-analysis to identify genes under positive selection. We thereby identify an inner membrane transporter (*ynjC*) with a putative role in colistin resistance. Contextual data from other sources can thus enhance local genomic surveillance at multiple levels and should be integrated by default when available.

## Introduction

Multiresistant strains of *Klebsiella pneumoniae* (Kp) are a global health threat.^1^ Among all known resistance mechanisms, carbapenemases are the most concerning, as they render most clinically relevant antibiotics ineffective.^2^ These enzymes are typically encoded on mobile genetic elements such as the Tn4401 transposon,^3^ which mediates transfer between plasmids^4^ and bacterial species.^5^ Furthermore, the prevalence of carbapenemase-producing Kp has increased in recent years.^6^ Such pathogen spread can be prevented by molecular surveillance and derived public health measures: Isolate genomes reveal transmission routes by accumulating characteristic mutations, from which ancestry can be inferred through time-dated phylogeny.^7^

While it has become standard practice to reconstruct such phylogenies of within-hospitasl outbreaks,^6,8^ few studies assess “contextual” information, i.e., genome sequences from isolates that were not part of the local outbreak but closely related. From a public health perspective, this is suboptimal. While many larger hospitals run screening programs to detect the carriage of resistant strains on admission,^9,10^ peripheral institutions rarely do. However, there is a significant transfer of patients, e.g., from operation theater to rehabilitation center or from one country to another. For an outbreak investigation with only local scope, these boundary-crossing transmission events remain hidden.

We here reanalyze a large outbreak at the University Hospital Leipzig (UHL) from 2010-2013^11^ in light of new data from a nearby institution, which experienced an outbreak with a closely related, albeit non-descending, strain. We performed a genomic meta-analysis to link both outbreaks, discovering hundreds of related genomes distributed across 28 different studies. We identify the likely sources of both outbreaks and illustrate hidden transmission across study boundaries. Only the integration of data from several sources provided a “complete picture”. However, we highlight several obstacles that need to be addressed before cross-boundary genomic surveillance can work in practice.

Beyond epidemiology, we show how outbreak meta-analyses can generate new hypotheses about host adaptation and antimicrobial resistance: The genomes under study underly similar selective pressures, such as treatment with colistin, an antibiotic of last resort. Thus, recurring mutations in the same gene(s) but across different genomes can signal putative causes for an observed phenotype, such as colistin resistance.^12^ For colistin, several such inducible genomic changes have been described that mediate resistance.^13^ Nevertheless, the exact mechanisms remain incompletely understood and seem to be multifactorial.^14^ We show how contextual data can be leveraged to generate hypotheses about putative factors contributing to colistin resistance.

## Results

### *In silico* and PCR-based screening identifies hundreds of outbreak-related, contextual genomes

Usually, hospital outbreaks are analyzed in isolation. However, it can be valuable to place local data in a larger genomic context. Such a context can inform about the origin and distribution of the outbreak-causing strain and reveal transmission routes. This knowledge then enables an effective public health response. From 2010 to 2013, UHL experienced a large outbreak of a multiresistant, *bla*_KPC-2_ -carrying Kp strain (hereafter referred to as “Kp-1”) of sequence type ST258, characterized by capsule type KL106 and O antigen (lipopolysaccharide, LPS) serotype O2v2. 105 patients were affected, and it took a multidisciplinary team many months to contain it^11^ (Figure S1). When we obtained 13 isolates from a 2018 outbreak of a *bla*_KPC-2_ Kp strain in a hospital nearby (“Kp-2”), we hypothesized that this strain was related to the previous outbreak at UHL due to its proximity in space and time.

A comparison of two genome sequences from Kp-1 and Kp-2, isolated from the respective index cases, showed that they were closely related, differing in only 69 single nucleotide variants (SNVs). While within-hospital Kp outbreaks have been estimated to differ at fewer than 21 SNVs,^6^ we are unaware of recommendations for isolates further apart in space and time. Therefore, more “contextual” genomes were needed to populate the genomic distance between Kp-1 and 2 and to fill the genomic “gap”. We, therefore, performed a comprehensive, multi-modal screening, consisting of (1) a comprehensive literature search including manual extraction of genomes and metadata, (2) an *in vitro* screening of two culture collections, and (3) an *in silico* screening of publicly available genomic and metagenomic datasets.

In total, we obtained 9,409 Kp genomes. Of those, 142 were collected during the Kp-1 outbreak from 105 patients,^11^ and 28 resequenced in parallel using long reads (Nanopore) to obtain accurate plasmid reconstructions. A further ten isolates were identified in two culture collections through PCR-based screening using strain-specific primers (see methods). Sequencing of the ten isolates confirmed that all were closely related to Kp-1. The primers were designed using a proprietary algorithm (nanozoo GmbH) to recognize Kp-1 and close relatives but not other Kp strain genomes, e.g., different sequence types. Interestingly, the algorithm selected a putative intact prophage region as the most specific PCR template, and all but two of the 415 total analyzed genomes contained the target (see methods). While phages are often considered mobile elements, they can be remarkably stable across decades.^15^

The remaining 9,257 genomes were collected from public sources. The majority was retrieved from NCBI *RefSeq*.^16^ However, 80 datasets were only identified through a literature survey, as they did not have an associated genome assembly deposited. In total, 28 studies were identified spanning 16 countries (Figure 1A and B, Table S1). In addition, extensive metadata were extracted where available. Furthermore, we searched the index Kp-1 isolate in a k-mer database of over 400,000 datasets of unassembled short reads (SRA, NCBI). We identified a single sample from an unpublished study of ICU patient colonization (SRA, project ID PRJNA561398) where we could recover a closely related, metagenome-assembled Kp genome.

**Figure 1:**
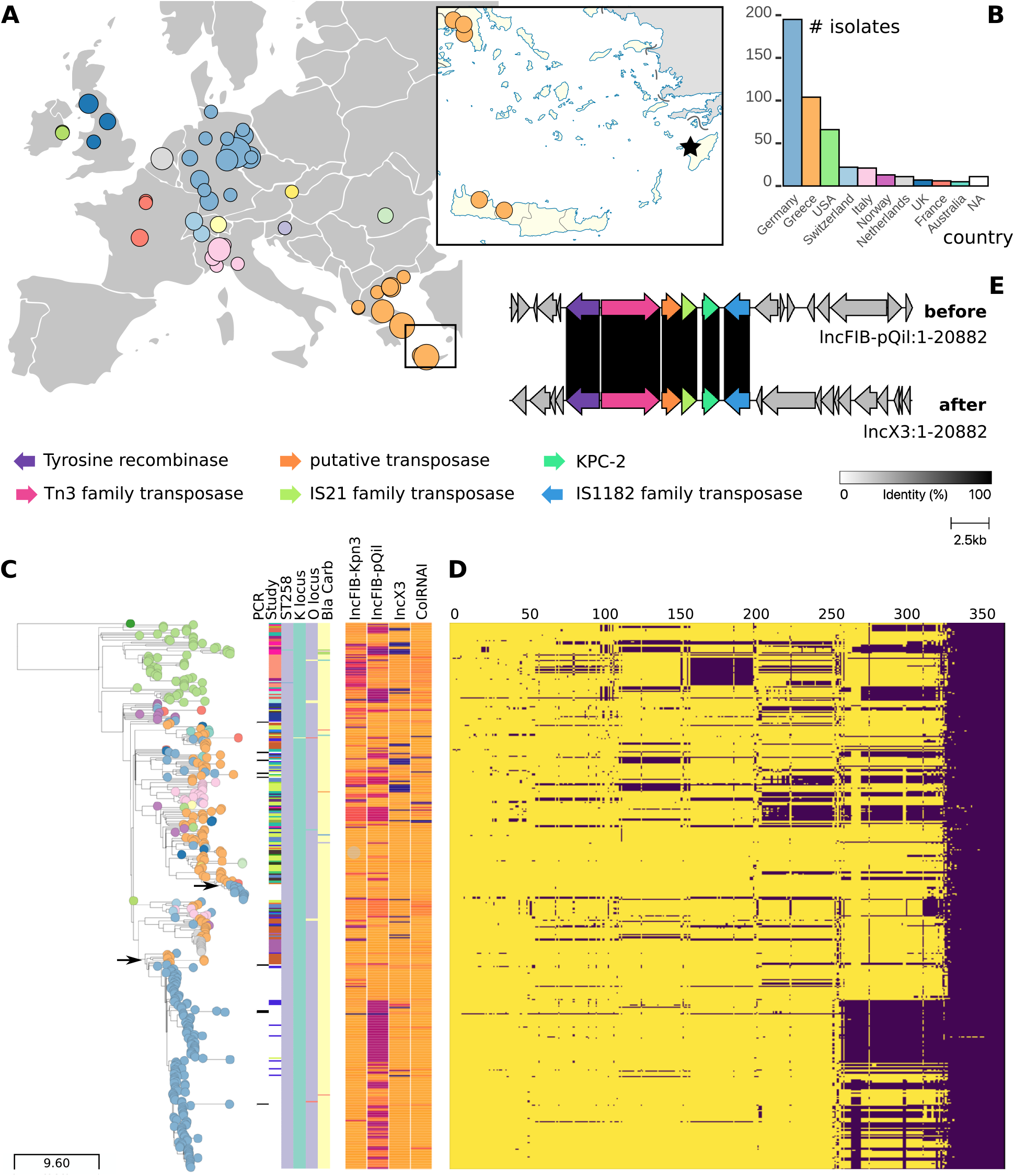
Time-dated phylogeny of 415 *bla*_KPC-2_ carrying Kp genomes and associated metadata blocks. Leaves colored by country code: Germany blue, Greece orange; for interactive exploration see visualization in microreact^25^ (microreact.org) under project ID 6bBfAYXswvY691LfbVytLT. **(A)** Geographic distribution of the genomes under study. Circle size is proportional to the number of genomes collected from this location. In the detailed map to the right, genomes found in Crete are shown (bottom). Our index patient was hospitalized in nearby Rhodes (star), and endemic transmission across these islands, which are connected by boat, is plausible. **(B)** Distribution of countries from which isolates were collected. **(C)** A timetree reveals how both outbreak strains, Kp-1 (lower arrow) and Kp-2 (upper arrow), most likely originated from southern and northern Greece, respectively (orange leaves above the arrows). In the leftmost metadata block columns, read from left to right, genomes are marked that have been identified using our strain-specific screening PCR. The 2nd column indicates which study they were recruited from (white is our study, all other 28 colors are study-specific, see Table S1). The next four columns show the sequence type (purple is ST258), capsule type (turquoise is KL106), O antigen (LPS) type (purple is O2v2), and carbapenemase variant (yellow is *bla*_KPC-2_). Note how the majority of genomes that pass our tiered filtering approach are homogenous in these features. The following metadata block shows plasmid containment as a fraction between 0 (blue) and 1 (orange) for four plasmids found in the index patient of Kp-1. **(D)** Matrix indicates presence (yellow) or absence (purple) of genes (columns) for each genome (row) in the phylogenetic tree. **(E)** Alignment of genes around the *bla*_KPC-2_ locus between the two plasmids IncFIB(pQil) (top) and IncX3 (bottom) shows a recombination event that allows shedding of IncFIB(pQil) while maintaining *bla*_KPC-2_ on the IncX3 plasmid, likely increasing host fitness.

Of the collected 9,409 genomes, 415 genomes (4.4 %) passed a tiered quality control protocol (see methods), resulting in a collection of high-quality genomes (ANI > 99.98 %, alignment to Kp-1 index isolate > 90 %) for further analyses.

### Time-dated phylogeny resolves outbreak origin and reveals hidden transmission

We observed 69 SNVs between the genomes of Kp-1 and Kp-2. Given the interval of seven years between the two respective isolation dates and a genome size of 5.3 Mb, this would correspond to a mutation rate of 1.85 per Mb per year, were Kp-2 a descendant of Kp-1. Because mutation rates of up to 1.42 per Mb per year have been reported in the literature,^17–19^ a direct relationship between both outbreaks seemed possible. We, therefore, constructed a time-dated phylogeny based on an alignment of 3,720 core SNV sites (total alignment length 5,384,856 sites) from the 415 genomes in our filtered collection to investigate these claims (Figure 1C). With it, we estimate the mutation rate of the corresponding Kp strain (ST258) to be 0.68 mutations per Mb per year (root-to-tip regression, *R*^2^ = 0.34). At this mutation rate, we expect each genome to experience one mutation about every 101 days (mean waiting time *t*)^20^ corresponding to 25 SNVs between Kp-1 and Kp-2 were they directly related, which is less than half of the distance observed. We, therefore, conclude that Kp-2 is not a direct descendant of Kp-1, which is supported by the reconstructed phylogeny (Figure 1C). The tree topology did not change when we used the mutation rates from the literature as fixed parameters in its construction.

The index patient’s travel history and symptom onset led to the hypothesis that the origin of the Kp-1 outbreak was a Kp strain imported from Rhodes, an island in southern Greece and a popular tourist location for German travelers. After being acutely hospitalized there, the patient was transferred to UHL, where *bla*_KPC-2_ was detected for the first time in the patient’s medical history. However, while a Greek origin seemed plausible, given the high prevalence of carbapenemase-carrying strains in this country,^21^ it could not be substantiated with data.^11^ In support of this view, we identified several closely related genomes from Crete,^22^ a neighboring island of Rhodes (see detailed map in Figure 1A), which populate the timetree around the time of the start of the Kp-1 outbreak (Figure 1C, lower arrow). With frequent travel by boat between these islands, it is plausible that an ancestor of Kp-1 was circulating in this region. Interestingly, the originating strain for the Kp-2 outbreak also seems to have come from Greece, albeit from northern provinces. Here, we could identify closely related genomes from two studies^6,23^ (Figure 1C, upper arrow). We even identified a third transmission from Greece to mainland Europe, with a strain from northern Greece causing an outbreak in the Netherlands^24^ (Figure 1C, grey leaves). The authors of the corresponding study did not identify this origin because they limited their investigation to local cases, supporting our argument for an integrative approach across study borders. All nodes in the tree where these transmissions out of Greece appeared had over 95% bootstrap support. However, it is important to consider potential sampling bias when inferring origins. While we identified many samples from Greece (Figure 1B), the screening methods were blind towards genome origin and considered an exhaustive set of Kp genomes. Furthermore, several studies have described the high prevalence of carbapenemase-carrying Kp in southern Europe.^21^ Therefore, we conclude that the large number of Greek samples likely represents the true distribution of *bla*_KPC-2_ Kp and is not an artifact of sampling bias.

As the Kp-1 outbreak unfolded, local health authorities assumed that the outbreak was likely not limited to one hospital. They based their assessment on the long duration and the large number of patients involved in the outbreak, with frequent transfers to and from the hospital as a tertiary care center. While these factors make non-local transmission more likely, no evidence was available to support this hypothesis. Surprisingly, we identified 13 isolates that were collected outside of UHL, but are part of the Kp-1 outbreak (Figure 1C, Table S2). Most of them come from the same federal state that UHL is in, but several were isolated in other states hundreds of kilometers away. No other countries were affected by the Kp-1 outbreak. The Kp-2 outbreak seems to have been contained within the affected hospital, as no published genomes were found in other places. On an international level, the data supports repeated introduction of KPC-carrying Kp strains from Greece, likely due to it being a popular travel site. In fact, travel-related carbapenemase-producing Enterobacterales have been recognized as an important source of resistance transmission.^26^ The above described hidden transmission events would not have been observed without integration of data across study borders, and illustrate the value of our approach.

### Carbapenemase preservation under frequent plasmid changes

Plasmids serve many functions, but a central one is as a gene delivery platform.^27^ Their payload is manifold, and here includes the *bla*_KPC-2_ carbapenemase. However, to the host genome, plasmids come at a considerable fitness cost, which creates pressure to remove them unless they provide a selection advantage.^27^ At the same time, plasmids resist removal through, e.g., toxin-antitoxin systems and compete with rivaling plasmids.^27^ In search of persistence in the host, frequent changes to the genetic material of plasmids can be observed.^28^ In the Kp-1 outbreak, we found four types of circular plasmid using Nanopore-based hybrid assembly: IncFIB(Kpn3), IncFIB(pQil), IncX3, and ColRNAl. We first quantified which fraction of each plasmid type from the Kp-1 index isolate was contained in all other genomes in our collection (Figure 1C). To complement this data, we aggregated the plasmid-encoded gene content (“pangenome”) across 28 Nanopore-sequenced isolates (Figure 1D). Note that the same gene can be carried by more than one plasmid in the same isolate. We find that genes on plasmids of type IncFIB(pQil) and less so IncX3 are frequently lost across our collection, illustrating the evolutionary forces described above.

Short read-based plasmid assemblies are often fragmented and incomplete and could mislead analyses. However, the Kp-1 outbreak allows for a more detailed study of plasmid dynamics, as we sampled 142 isolates and, more importantly, were able to reconstruct complete circular plasmids using long-read sequencing for 28 of them. We observed the complete loss of IncFIB(pQil) during the outbreak. This was initially confusing because this plasmid carried the *bla*_KPC-2_ carbapenemase, which was detected using culture on screening agar followed by qPCR in all isolates. We then found one isolate with two *bla*_KPC-2_ copies, one on IncFIB(pQil) and IncX3, respectively. Because *bla*_KPC-2_ is surrounded by transposases (Figure 1E), it readily recombines, and *bla*_KPC-2_ was copied from IncFIB(pQil) to IncX3. Thereafter, the host could discard the IncFIB(pQil) plasmid but retain the selective advantage of *bla*_KPC-2_. As indicated by the contextual genomes, IncFIB(pQil) loss is frequent and likely confers a fitness advantage. In line with this argument, all descendants of the isolate with two *bla*_KPC-2_ copies have discarded the IncFIB(pQil) plasmid and carry *bla*_KPC-2_ on IncX3 (Figure 1C).

### Contextual genomes reveal positive selection of virulence and resistance genes

Comparative genomics can reveal adaptations to specific stimuli under selective pressure, such as antibiotic treatment.^29^ The pathogens in our curated dataset can be assumed to be under similar selective pressures. For example, all but one were isolated from patients in hospitals (one isolate found in the literature was sourced from a wastewater plant^30^). Furthermore, since all included Kp isolates carry a carbapenemase, few antibiotics remain as a rational treatment option. One of them is colistin, sometimes combined with rifampicin for synergy.^31^ Additionally, the Kp isolates will likely evolve to facilitate, e.g., long-term carriage and virulence. Such adaptations can be detected when aggregating mutations for each gene across all genomes:^32^ If the rate of non-synonymous substitutions (*dN*) is higher than the rate of synonymous substitutions (*dS*), positive selection of the affected gene is plausible.^33^ This signal can, in turn, generate new hypotheses about the gene’s function.

Modified lipopolysaccharides often cause colistin and, more generally, polymyxin resistance (PR). They result in a positive charge to the bacterial membrane that repels polymyxins.^13^ Several proteins are involved, though *“the exact mode of action of polymyxins still remains unclear*.*”*.^13^ For 171 of the 415 genomes in our collection (39.7 %), we were able to assess from the original publications whether the isolate was colistin-resistant or not (Table S2). Where minimum inhibitory concentration (MIC) measurements were available, breakpoints by the EUCAST committee (v11) were used to classify isolates into colistin sensitive and resistant. To identify genomic regions associated with PR, we first performed a genome-wide association study (GWAS) based on SNVs, including small insertions and deletions. This analysis did not return a significant result after correcting for population structure (*p* > 0.05), neither when considering each SNV individually nor when aggregating SNVs over genes in a so-called *burden test*.^34^

This failure might be due to technical limitations of GWAS, especially in light of few genomes^35^ or strong population structure.^36,37^ Furthermore, while single SNVs can induce colistin resistance,^38^ PR is generally assumed to be a polygenic phenomenon.^13^ To test which genes were mutated more than expected, we first aggregated unique haplotypes for each gene across all genomes, similar to a *burden test*.^34^ To conservatively correct for population structure, we counted mutations only once per position in the reference genome. Recombinant sites, putative phages, and sites within repetitive sequences were excluded. This procedure would not detect convergent evolution where mutations arise in the same position in two different clades. However, we did not detect any homoplastic mutations outside of recombinant regions. At a mutation rate of 0.68 per Mb per year, over the study period of our meta-analysis of about ten years, and assuming 4,000 genes per genome, we expect the number of mutations per gene to follow a Poisson distribution with a mean of 0.01 mutations per gene. This estimate is supported by the data where most genes remain unchanged (Figure 2B).

**Figure 2:**
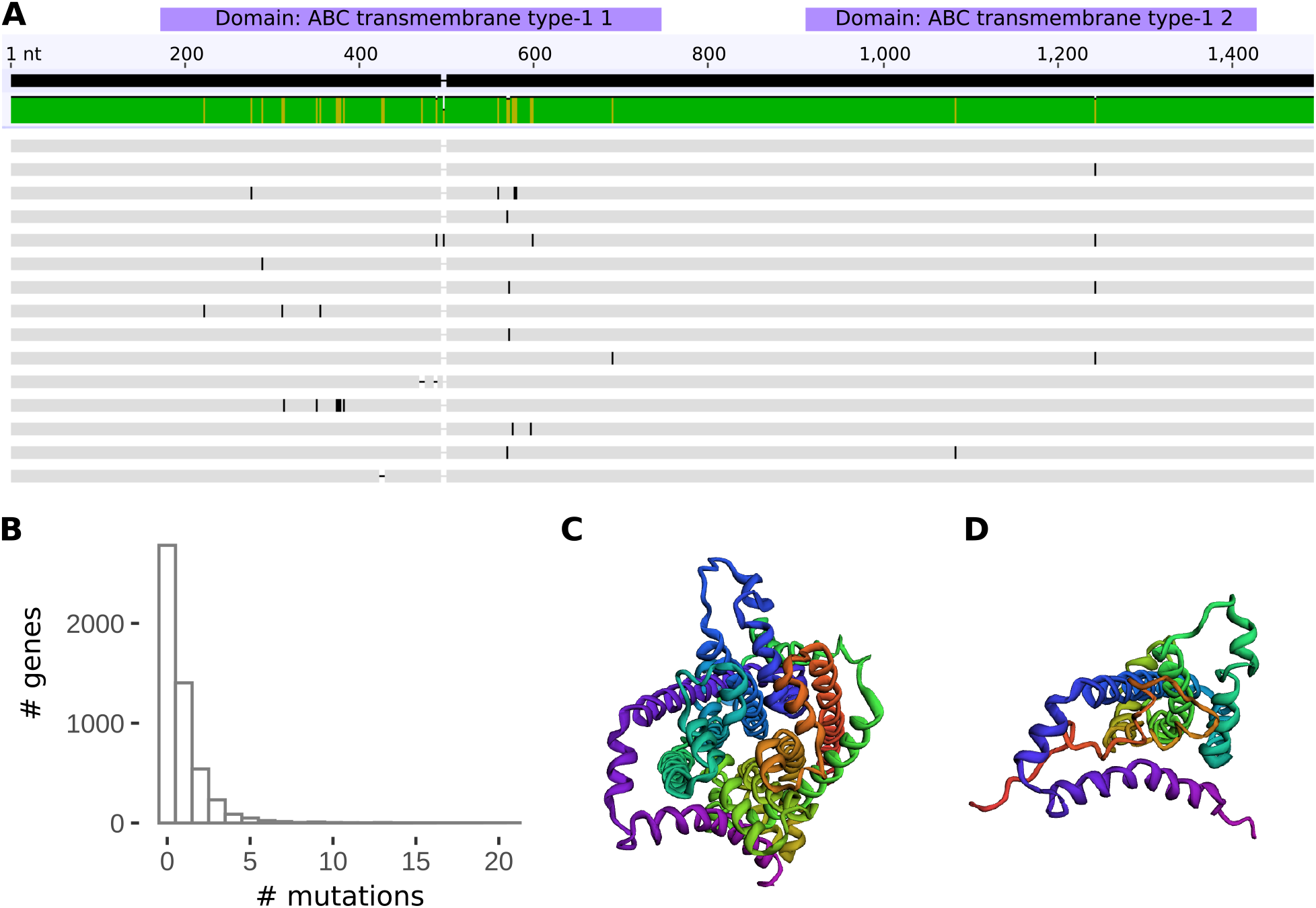
Positive selection of the inner membrane ABC transporter permease *ynjC*. **(A)** Multiple sequence alignment of representative haplotypes of the nucleotide sequence of *ynjC*. Most mutations occur between positions 200 to 700, which includes both transmembrane and interacting domains. Three of those haplotypes lead to premature stop codons. **(B)** Distribution of unique mutations observed in all genes. As expected by the estimated mutation rate of 0.68 mutations per Mb per year, most genes remain unchanged over the ten years which our study covers. Several genes, however, accrue over 20 unique mutations across 415 genomes. **(C)** 3D protein structure of the *ynjC* permease. In the center is the pore through which small molecules are shuttled. **(D)** 3D protein structure of a truncated form of the protein (same orientation as C), created through a premature stop codon. Clearly, the channel structure is lost, and the protein is likely dysfunctional.

We ranked genes by the number of unique mutations (UM) per gene. We define unique mutations as specific to a single genome and position, and we used the number of UMs per gene as a heuristic to rank and prioritize genes for further analyses. A gene set enrichment of all genes with ≥ 10 UMs (n=15) showed two overrepresented biological processes. For one, the phosphorelay signal transduction system was enriched (19.8-fold, *p* ≤ 0.001),^39^ which is known to be implicated in PR.^13^ Furthermore, genes associated with nitrate assimilation were enriched (41.8-fold, *p* ≤ 0.001), which to our knowledge has neither been described nor could we assess the biological significance of this finding.

We tested the top-ranking genes with the highest number of unique mutations for gene-wide evidence of episodic positive selection. For each candidate gene, we used a random-effects framework to pool evidence across multiple sites and thereby increase statistical power.^32^ All genes discussed hereafter exhibited significant positive selection (*dN/dS* > 1, likelihood-ratio test, *p* ≤ 0.05). We found two positively selected genes that affect virulence: The transcriptional activator *cadC* (18 UM) has been linked to increased Kp colonisation^40^ and *fimH* (16 UM) is a critical virulence factor in urinary tract infection, a common complication of Kp colonisation.^41^

It is plausible to assume that most isolates in our study are subject to similar treatment-associated adaptive pressure: Since most beta-lactam antibiotics fail to treat these isolates, colistin will have been used in many patients. We found several known genes involved in PR to be mutated. In 47 out of 171 isolates with an available phenotype (27,5 %) we found a truncated or missing *mgrB* gene product, a negative regulator of the *PhoPQ* signalling system.^42^ 39 of these 47 (83 %) were resistant to colistin. This adaptation can occur rapidly: In the single Kp-1 outbreak, we identified three different *mgrB* loss-of-function mutations (Figure S2). Furthermore, we found frequent truncations in *pmrB* and non-synonymous mutations in *phoQ* (17 UM), both regulatory proteins involved in LPS modification.^43^ These “canonical genes”^42^ cause PR by acting on the outer membrane. We did not detect the plasmid-encoded *mgr-1* gene, which encodes a transferase that modifies lipid A and thereby causes PR.^13^

Recently, colistin has also been found to target the inner cytoplasmic membrane.^44^ Interestingly, we identified a highly mutated inner membrane ABC transporter permease^45^ under strong positive selection (all detected mutations non-synonymous), named *ynjC* (21 UM, Uniprot, P76224). Proteins of this group utilize ATP to import many small molecules such as nutrients and antibiotics.^46–48^ Mutations in permeases have been shown to “lock” the transporter in one of its two states,^49,50^ such as inward-facing,^51^ disrupting the shuttle function.^52^ Additionally, we found three mutations that caused premature stop codons and subsequent dysfunctional proteins (Figure 2C and D). Most mutations accumulate in a region between residues 75-230, spanning both transmembrane and topological domains (Figure 2A). In 12 isolates with *ynjC* mutations, 7 (58.3 %) were resistant to colistin; however, for none of the haplotypes with premature stop codons, phenotype data could be obtained, and future functional validation is needed. However, ABC family transporters have been proposed to transport nascent core-lipid A molecules across the inner membrane,^53^ with a putative effect on colistin resistance. They have also been proposed as an antibiotic target.^54^ We thus argue that the *ynjC* permease could have a role in PR.

## Discussion

Genomic surveillance is a powerful public health tool to reduce the spread of resistant bacteria. We show that genomic meta-analysis of outbreak genomes can provide important contextual information when interpreting local outbreaks. To construct the context, we employed both *in vitro* and *in silico* search methods to aggregate more than 400 genomes to supplement the local outbreak under investigation, screening more than ten thousand genomes and half a million metagenomes in the process. As a result, we discovered critical epidemiologic details that would have been missed in a traditional outbreak study focusing on local data only. For example, we determined the likely source of the Kp-1 outbreak, its relation to an outbreak at a nearby institution, and it being an instance of the repeated introduction of *bla*_KPC-2_ Kp isolates into mainland Europe from Greece. We also identified isolates from other studies that are direct descendants of Kp-1.

We then illustrated the plasmid dynamics across our genome collection. We found frequent loss of genetic material associated with IncFIB(pQil)-type plasmids, even though they often carry the *bla*_KPC-2_ gene. We resolved this paradox by showing how *bla*_KPC-2_ can still be preserved in the host: The carrier transposon is first transferred to another plasmid before IncFIB(pQil) removal from the host.

Besides phylogenomic insights, our context-enriched genome collection informs about adaptation to selective pressure. For one, we found several positively selected genes that are known to mediate, e.g., colistin resistance. We also discovered positive selection of the inner membrane transporter *ynjC* together with an overrepresentation of mutated gene copies in colistin resistant isolates. However, future experiments will have to validate if an effect on colistin resistance can indeed be shown, e.g., by introducing loss-of-function mutations using CRISPR.^42^

Several components are still missing until we can analyse putative outbreak genomes in a real-time, integrated surveillance system. The main bottleneck, counter-intuitively, is not sequencing but data management and bioinformatics.^55^ For example, there is no common repository for bacterial outbreak metadata in active use by the community. We manually aggregated metadata from 28 studies, which frequently involved squinting at low-resolution images to extract, e.g., data on colistin resistance. For most genomes, important information besides the year and country of isolation was missing. Without this metadata, the sequenced genomes cannot easily be integrated into any analysis other than the one they were originally sequenced for. This could be aided in the short term if authors published supplementary data giving genome accessions alongside all relevant isolate data, genotypes and phenotypes explored in the study.

Also, more sophisticated tools for outbreak genome sharing are needed:^56^ Most outbreak studies appear one to two years after the outbreak took place (personal observation). However, by then, the value of the results is primarily academic. Only prospective data analysis^57^ in real-time would enable a practical outbreak response. A recent example of this is `nextstrain`, where the virus genomics community converged on a set of protocols and databases,^58^ which allowed a data-driven public health response. When combined with real-time sequencing of bacterial genomes,^59^ this set of technologies could substantially improve outbreak response.

## Methods

### Culture and Sequencing

142 Kp-1 isolates were collected from 105 patients in a previous investigation^11^ and complemented in the present study with an additional ten isolates discovered using PCR screening of two culture collections (see below).^11^ 13 samples were collected from Kp-2. All of the isolates were sequenced using short reads (Illumina). 28 Kp-1 samples were additionally sequenced using long reads (Nanopore) to enable hybrid assembly (see below). All samples were streaked on CHROMagar KPC chromogenic agar plates (CHROMagar, Paris, France), and KPC carriage was confirmed using PCR. DNA extraction for Nanopore sequencing and quality control was done as reported elsewhere.^60^ Care must be taken, especially for Nanopore sequencing, not to damage the extracted DNA to achieve a sizeable median fragment length (target 8 kb) for sequencing to be effective. Nanopore sequencing was performed using the MinION sequencer and the 1D ligation library kit (LSK109) on an R9.4 flow cell (all Oxford Nanopore Technologies, ONT). Illumina sequencing for isolates from other studies is described in the respective publications (Table S1). For genomes resequenced for the current study, a read length of 150 bases (paired-end) was used on an Illumina MiSeq sequencer. The libraries were constructed using a previously established protocol.^61^

### In silico screening of isolate and metagenomes

In screening, our aim was to collect as many genomes as possible with a putative relation to the outbreak clone Kp-1, yielding a total of 9,409 genomes. From NCBI *RefSeq*, we retrieved all 9,177 genomes that were labelled as *Klebsiella pneumoniae* (Taxonomy ID: 573, last access 2020-08-01).^16^ In a comprehensive literature search using the search terms “KPC, Klebsiella pneumoniae, outbreak” we identified 80 genomes from various studies that had only deposited reads with NCBI SRA, and which we reassembled for this study (see below).

For metagenomic search, we screened about 500,000 metagenomic read sets in a reduced representation known as *MinHash* signature^62^ using wort (no version, unpublished, github.com/dib-lab/wort). Hashing was performed using sourmash (v3.5).^63^ As query we used the Kp-1 index genome (k=51, sampling rate 0.001) and manually reviewed all 15 hits reported with a threshold ≥ 0.01 Jaccard similarity, a measure that approximates average nucleotide identity (ANI).^62^

### Strain-specific screening PCR

We then screened two culture collections (National Reference Center for multidrug-resistant Gram-negative bacteria, Bochum, and Medical Microbiology and Virology, Leipzig) for related isolates using a strain-specific marker PCR, designed using a proprietary, pangenome-based algorithm (nanozoo GmbH). Each 50 µL PCR reaction contained 10 µL template DNA, 2 µL 10 nM primer mix for each primer (primer 1: ATGCGTCCACGAAGAATTAT, primer 2: CATCGCCAAGATACTGTACA), 25 µL 2x polymerase master mix (Superfi II, Invitrogen) and 11 µL ultra-pure water. Thermal cycling consisted of initial denaturation at 98 °C for 1 minute followed by 35 cycles of denaturation at 98 °C for 20 s, annealing at 55 °C for 20 s, extension at 72 °C for 1 min, followed by final extension at 72 °C for 5 min.

### Data processing

Unless otherwise stated, default parameters were used. Of the 9,413 collected genomes, 1,461 passed a minimum Jaccard similarity of 0.97 (15.5 %, parameters: k-mer size 51 nt, scale 0.001). Jaccard similarity was computed using sourmash (see above). In a subsequent filtering step, 415 (4.4 %) were included for tree construction based on a minimum *in silico* DNA-DNA-hybridization threshold of 99.98 % computed using FastANI (v1.32)^64^ as well as a minimum genome length of 5 Mb and an alignment of 90% of the query genome to the Kp-1 index isolate (completed, circular), excluding all extra-chromosomal sequences. This sequential approach allows for laxer but computationally efficient methods with fewer constraints to screen many genomes in the beginning. Subsequently, the selection is refined using more computationally expensive methods. We conservatively removed 16 samples from the timetree because they did not fit the estimated molecular clock model, likely due to unidentified recombination.

Isolates where only short reads could be obtained were assembled using shovill (v1.1.0, unpublished, github.com/tseemann/shovill). Metagenomic reads were preprocessed using fastp (v0.20.1)^65^ and assembled using megahit (v1.2.9).^66^ All contigs with a minimum length of 2 kb were then mapped to the reference genome (Kp-1 index patient, VA13414, Table S1) using minimap2 (v2.17-r941)^67^ with the asm5 option for an expected sequence divergence of ≤ 0.1%. The Nanopore sequencing data were basecalled using Albacore (v2.3.2, available from Oxford Nanopore Technologies) and adapters removed using Porechop (v0.2.3, unpublished, github.com/rrwick/Porechop). Genome hybrid assembly using long and short reads was performed using Unicycler (v0.4.6).^68^

Genome annotation was performed using prokka (v1.14.6).^69^ Annotation of Klebsiella-specific features was done using kleborate (v0.4.0-beta).^70^ Plasmids were annotated using abricate (v1.0.1, unpublished, github.com/tseemann/abricate) using the plasmidfinder database (version 2021-01-13).^71^ Antimicrobial resistance genes were annotated using the same program with the *Comprehensive Antibiotic Resistance Database* (CARD, v3.1.2).^72^ Phages were annotated using uv (v0.1, unpublished, github.com/phiweger/uv). Recombinant regions were annotated using gubbins (v2.4.1).^73^ SNV calling was performed using the snippy workflow (v4.6.0, unpublished, github.com/tseemann/snippy) which proved the most accurate program in a recent benchmark study.^8^ In short, snippy simulates reads from input genomes and maps them to the provided reference using bwa (v0.7.17-r1188),^74^ before calling variants with freebayes (v1.3.2, unpublished, github.com/freebayes/freebayes). Putative recombinant, repetitive and prophage regions were masked before SNV calling. Sites with SNVs were extacted using snp-sites (v2.5.1).^75^

### Reconstruction of time-dated phylogeny

A time-dated phylogeny was calculated using timetree (v0.7.6),^76^ a maximum likelihood-based approach starting from a core genome SNV alignment. The derived mutation rate was scaled by the total genome size. Homoplasy was assessed using the same approach. The final tree and associated metadata were visualized using the microreact webservice.^25^ Bootstrap support values were extracted from the guide tree, a prerequisite of the timetree, and calculated with raxml-ng (v0.9.0).^77^

### Analysis of genomic variants

A genome-wide association study was performed using pyseer (v1.3.7)^78^ and included the aggregation of mutations across genes in a *burden test*.^34^ Gene set enrichment was performed using the Gene Ontology webservice (last accessed 2021-04-01).^79,80^ Positive selection was assessed by first aligning all sequences for a particular gene using nextalign (no version, unpublished, github.com/nextstrain/nextclade). The multiple sequence alignment was then analyzed using the BUSTED algorithm^32^ as part of the HyPhy suite (v2.5.31).^81^ *In silico* folding of proteins was done using the trRosetta model (no version).^82^

## Appendix

## Ethical approval

This retrospective study was performed in accordance with the ethical guidelines of the 1964 Declaration of Helsinki and its later amendments and was approved by the local ethics committee (University of Leipzig, register no. 411-12-11032013). The need for informed consent was waived according to the ethics approval.

## Data availability

All data and metadata used in the analyses have been deposited with the *Open Science Foundation* (OSF) under project ID n78q3. Extensive metadata on all samples used in this study is available there and in the supplement (Table S1), including curated phenotype data on colistin resistance (Table S2). In addition, raw sequencing data generated under this study has been deposited with the *European Nucleotide Archive* (ENA) under project ID PRJEB45529. For all remaining raw data, please refer to the corresponding studies (Table S1).

## Author contributions

AV designed the study. AV, CBl and NL performed all laboratory work. AV and JH screened isolates using PCR. AV and CBl implemented Nanopore sequencing. AV, CBl and NL collected metadata. AV, CB, NL, KLW, CBr, LI and MH conducted data analysis. BG supervised the work. All authors interpreted the results, wrote the text, created the figures, and approved the submitted paper.

## Acknowledgments

The authors would like to thank Prof. A. C. Rodloff for extensive discussion of the manuscript. The authors would also like to thank T. Kaiser (Institute of Laboratory Medicine, Clinical Chemistry und Molecular Diagnostics, University Hospital Leipzig) and K. Finstermeier (Max Planck Unit for the Science of Pathogens, Berlin) for access to and guidance through the collected Kp-1 outbreak metadata.

## Supplement

**Figure S1:**
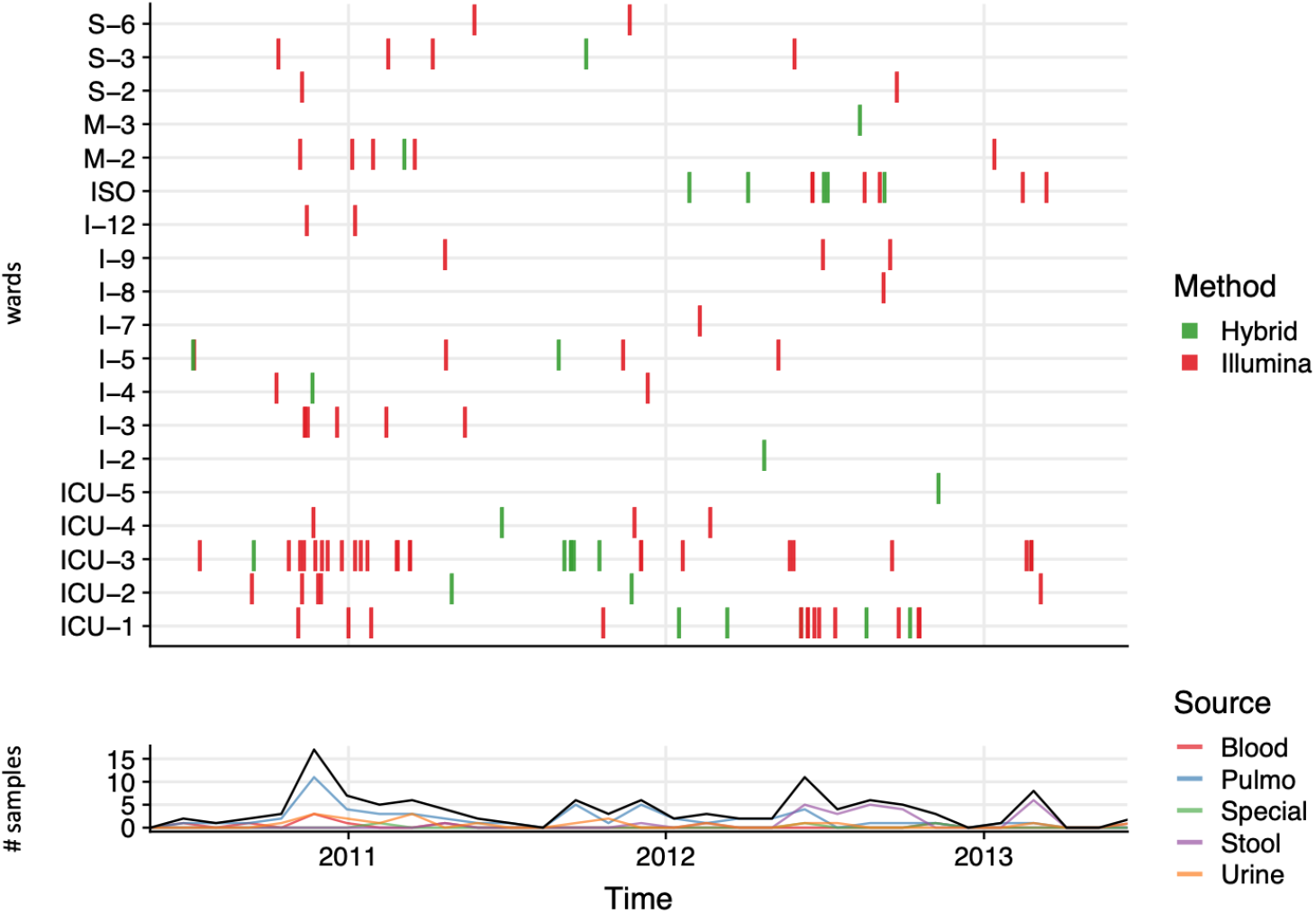
*bla*_KPC-2_ Kp-1 case distribution. All isolates were sequenced using short reads. For long-read sequencing, 28 representative samples were selected, uniformly distributed across time and space.

**Figure S2:**
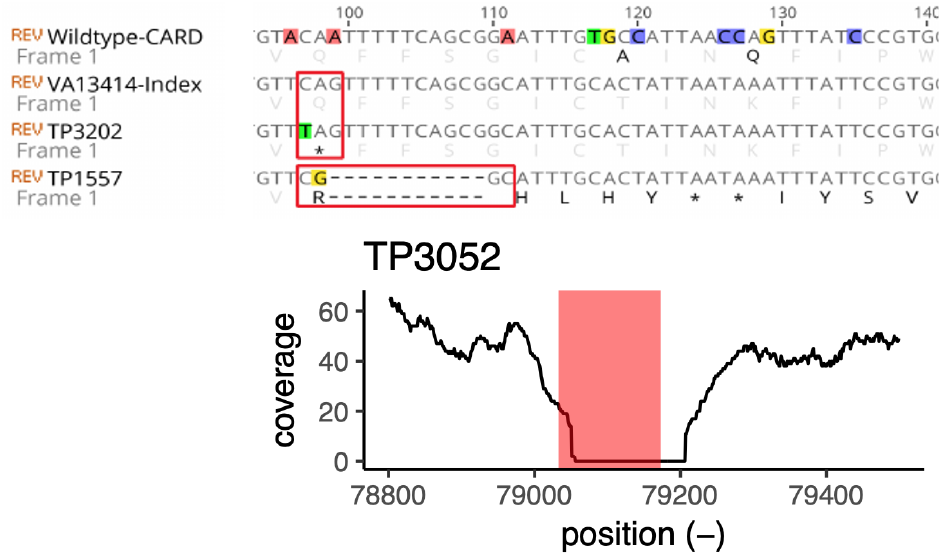
**(Top)** Detail from a multiple sequence alignment of representative *mgrB* sequences. Three different variants caused *mgrB* truncation in the Kp-1 outbreak, two of which are illustrated here. From top to bottom: Reference *mgrB* sequence from the CARD database, gene sequence from the Kp-1 index isolate, *mgrB* where a SNV causes a premature stop codon, gene sequence with an 11 bp deletion and subsequent frame-shift. **(Bottom)** One isolate presented a complete loss of *mgrB*, as could be validated by mapping the short reads from this isolate to the *mgrB* locus in the Kp-1 index genome.

